# Direct Measurement of Trunk Volume in Forest Trees: Focus on White Pine and Comparison to a Statistical Method

**DOI:** 10.1101/2020.03.18.995985

**Authors:** Robert T. Leverett, David N. Ruskin, Susan A. Masino

**Affiliations:** Native Tree Society (www.nativetreesociety.org), Senior Advisor to American Forests National Champion Tree Program, Washington, DC, USA; Friends of Mohawk Trail State Forest, Florence, MA, USA; Trinity College, Hartford, CT; Charles Bullard Fellow, Harvard Forest, Petersham, MA

**Keywords:** above-ground volume, cone, forest mensuration, form factor, frustum, neiloid, paraboloid, sine method, volume modeling

## Abstract

Accurate measurement of tree volume and associated carbon storage are necessary to determine ongoing sequestration as well as site productivity and changes in growth of individual tree species. Standard statistical methods vary their estimations of tree volume, and thus carbon storage and sequestration, particularly in larger, older trees in a forest setting. Here, we describe a detailed direct measurement method that combines traditional trunk taper models with state-of-the-art instrumentation and the best mathematical models for producing more accurate measurements of trunk volume. A stand-grown Eastern White Pine (*Pinus strobus*) is used as an example; the method is compared with a commonly used statistics-based Forest Service method. This latter method is shown to over- or underestimate volume if the trunk form factor deviates sufficiently from the average value for this species. Direct measurement modeling can be used to validate or choose among existing simple statistical volume models, especially for local applications. It can also assist in widespread recalibration of other standards and models used to estimate volume and carbon storage over time.

## Introduction

Trees yield products such as wood, food and medicine, along with nature-based services such as water and air filtration, and carbon storage and sequestration [1,2]. These and numerous other properties provide economic, environmental and social value, and many of these properties depend primarily on tree volume. For example, forests are the biggest source of negative carbon emissions, an essential factor in stabilizing the temperature and mitigating future climate change [4,5]. Therefore, tree volume, current carbon storage and potential carbon sequestration must be quantified accurately. Direct ground-truthing of tree volumes is essential for climate models, payment for ecosystem services, and forest management.

Two common problems that create inaccuracies in individual tree trunk volume determinations are 1) height-measuring errors, and 2) inaccurate diameter measurements at various heights going up a trunk. These initial inaccuracies in basic dimensions are magnified when calculating above-ground volume, which is then used to estimate below-ground volume. This report describes validation and application of calibrated traditional and state of the art instruments, and customized formulas to calculate the trunk volume of large trees accurately, and thus track changes over time. The method is illustrated using a stand-grown Eastern White Pine, and is compared to a common Forest Service Inventory Analysis statistical model in forty trees.

## Materials and Methods

Since 1990, the Native Tree Society (www.nativetreesociety.org), formerly the Eastern Native Tree Society, has been developing, refining and validating detailed field methods and protocols for measuring trees and calculating their volume. Equipment, procedures and calculations are provided.

### Equipment

#### Height Measurement

##### Laser Technology Inc (LTI) TruPulse 200X hypsometer

Used for most of the tree height measuring.

##### LTI Impulse 200LR hypsometer

Used as a periodic check on the TruPulse 200X. Although it is bulkier, it is more stable in its measurements at distances of 50 m or more.

#### Diameter and Limb Measurement

##### LTI TruPulse 360 hypsometer

Used to establish limb lengths. Its missing line routine can measure the straight-line distance between two points in three-dimensional space.

##### LTI TruPoint 300 mini-surveying station with special tripod adapter

Used to help gather diameter measurements of trunks, and also to use its missing line routine for crown spread.

##### Bushnell Legend Ultra HD 10×42 monocular with range-finding reticle

Used to measure diameter from a distance and at tree-heights above what can be reached.

##### Vortex Solo RT 8×36 monocular with range-finding reticle

Used to measure diameter from a distance and at tree-heights above what can be reached

##### 36-inch calipers

Used to measure trunk diameter near ground level and up to approximately 5.7 m.

##### Standard diameter tape (D-Tape)

Used to measure trunk diameter near ground level and up to approximately 5.7 m.

#### Accuracy Measurement

##### Bosch GLM80 and GLR825 construction lasers

Used to establish accuracy limits for the TruPulse 200X.

### Measurements

#### Tree and Frustum Height

Because the TruPulse 200X was the primary height-measuring device, its infrared pulse-based laser was repeatedly tested against class II phase-based lasers. The results of repeated tests of the TruPulse against a Bosch GLR825 construction laser yielded accuracies for the TruPulse within +/-2.5 cm to clear targets. The accuracy of the GLR825 is rated at +/-1.0 mm.

Data collected and verified show that the TruPulse 200X performs better than the manufacturer’s specifications (accuracy of +/-4.0 cm) to clear targets. Overall, it proved to have an accuracy of +/-2.5 cm to easily discernible targets, and the manufacturer value of +/-4.0 cm is probably most applicable to distant targets (such as leaves and stems in the canopy of a tree).

The Impulse 200LR can also measure tree-height. It is bulkier, and less likely to be used for practical reasons, but slightly more accurate than the TruPulse 200X. Accuracy tests show it to be accurate to +/- 1.5 cm.

#### Options for Measuring Height

The Tangent Method (Figure 1) is the traditional tree height measuring technique (e.g. [5]. It is implemented manually with a tape and clinometer, and automatically with hypsometers with a tree-height measuring function.

**Figure 1.**
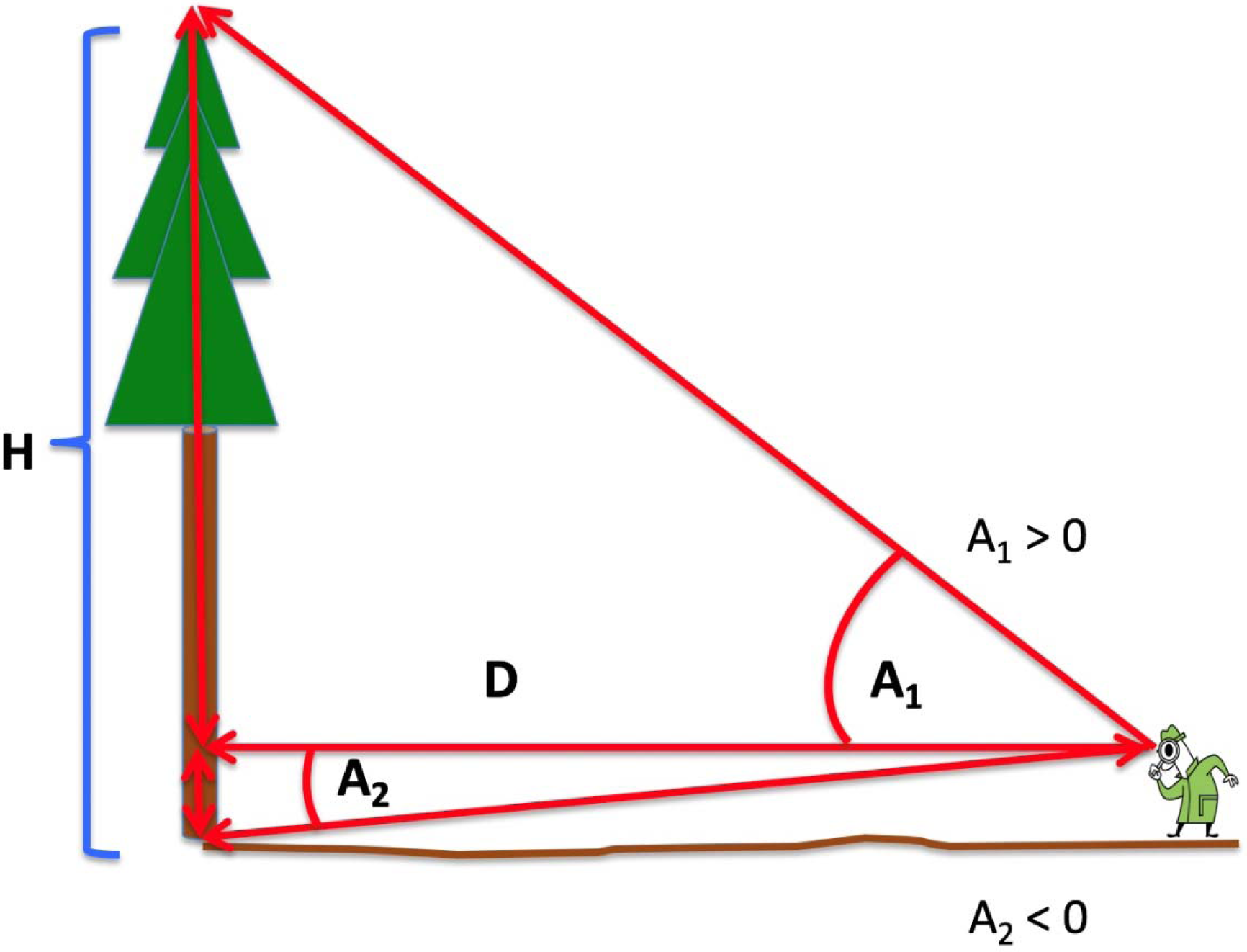
The Tangent Method of height determination illustrated. The distance to the tree at an arbitrary point on the trunk and the visible angles from this height to the base and top of the tree are measured. The method assumes that the top of the tree is directly over the base.

To implement the Tangent Method with a modern hypsometer, three separate shots are taken. First, the level distance to the trunk is measured, followed by angles to the base and to the top (in some cases what is chosen as the top, or direction of the top). Here is the formula for the Tangent Method:

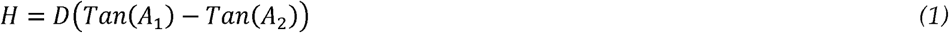

Where *H* = height

*D* = distance to 1.37 m from base (breast height)

*A*_1_ and *A*_2_ are angles as illustrated in Figure 1.

Angles below eye level are negative. For the Tangent Method to work as ordinarily used, and as shown in Figure 1, a tree’s top must be located vertically over its base. Therefore, the Tangent Method assumes that the base, the end of the baseline, and the top are in precise vertical alignment. If not, errors occur when the base line is taken level to the trunk. While vertical alignment is usually true for younger trees in a stand or for trees in plantations, strict vertical alignment is seldom the case for older trees that have lost apical dominance. In addition, the ends of up-turned branches horizontally offset from the trunk can be mistaken for the top when a tree becomes exceptionally tall and broad-crowned and the measurer is too close to the trunk. This situation commonly occurs in taller, older forests.

There are situations where the Tangent Method produces accurate results (younger tree, vertical alignment). It is valuable in cases when it is the only method that can be used, i.e. the top is not visible, but its direction can be reasonably estimated. However, it is usually possible to achieve better accuracy and avoid shortcomings of the Tangent Method by applying the Sine Method (Figure 2), a measurement technique that uses laser rangefinders and angle measurers (tilt sensors in hypsometers). The Sine Method depends on straight-line measurements from eye to target [6] and Figure 2 illustrates how the Sine Method allows the measurer to get the correct height regardless of where the top (assuming it is visible) is located within the crown. Angles below eye level are negative. Here is the formula for the Sine Method:

**Figure 2.**
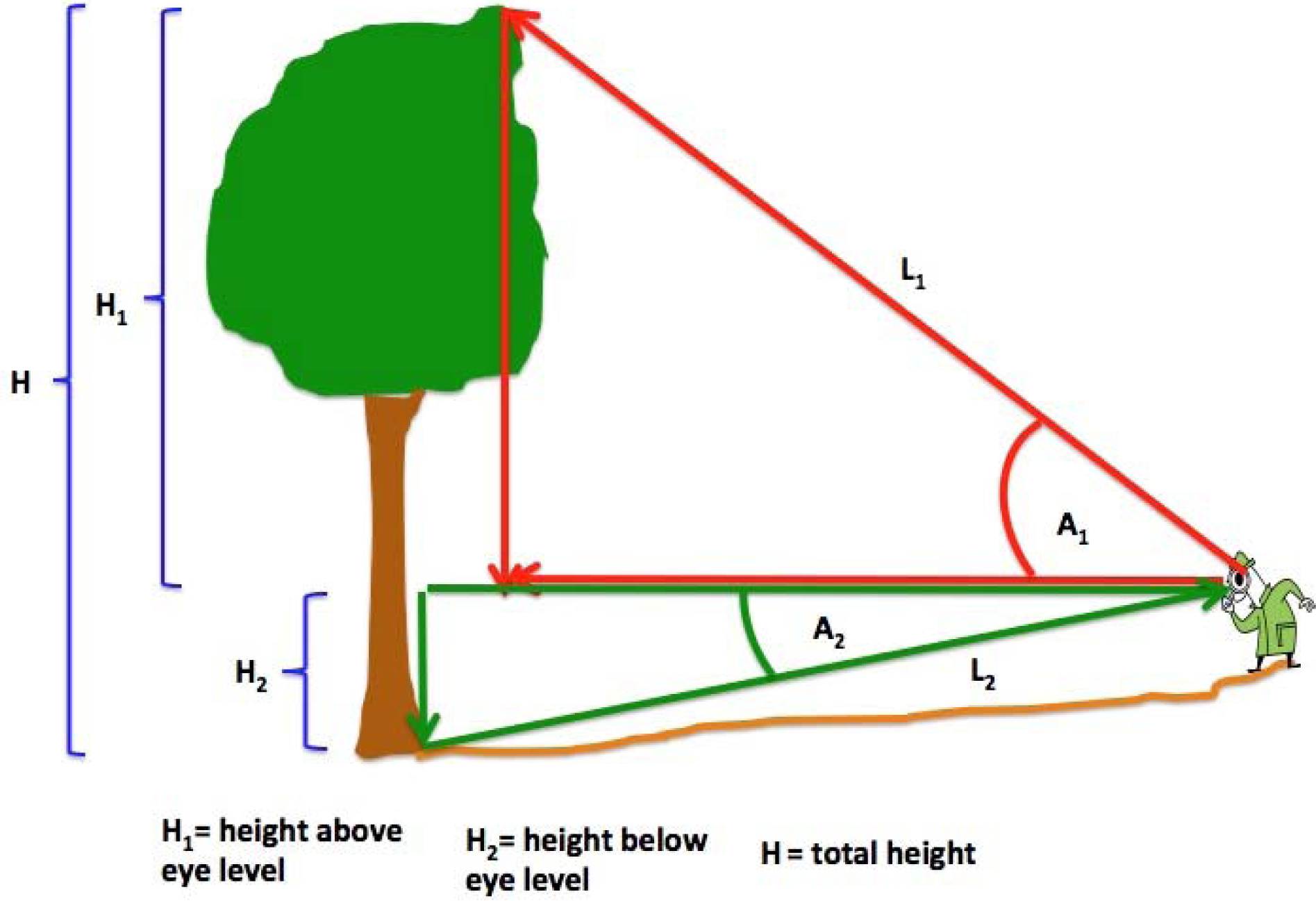
The Sine Method of height determination illustrated. The distances to the tree’s base and top are determined separately, as well as the visible angles from the observer’s position to the base and top. Verticality of the tree is not assumed.

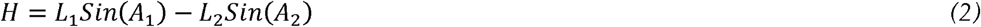

Where *H* = height

*L*_1_ and *L*_2_ are lengths to the top and the base, respectively, illustrated in Figure 2

*A*_1_ and *A*_2_ are angles as illustrated in Figure 2.

The Sine Method is inherently more accurate for the targets that are not vertically aligned, since distances are measured directly to the targets, and therefore, require an unobstructed sightline. Other than due to equipment error, the height of the target is not over-estimated, which is often the case with the Tangent Method. Here, if the measured spot in the crown is not the absolute top, the measurement will be less than the full height of the tree – but not more – and avoids over-estimating volume in older trees by inadvertently over- measuring height. There are methods for addressing a lack of verticality with the Tangent Methods; however, distance and angle errors have a greater impact on height with the Tangent Method than do the same errors incurred with the Sine Method, suggesting an overall superiority of the Sine Method. Regardless of method, the less the observer’s distance from the tree with an offset top, the greater the error in height measurement: one surveyor’s chain length (20.1 m, a commonly used distance) is inadequate for a tall mature tree with a broad crown.

Modern hypsometers often include routines for implementing both the Sine and Tangent Methods, with the Tangent Method often called the Tree-Height Routine. A manufacturer’s accuracy claims for their instrument should not be confused with the accuracy or appropriateness of the actual method used to measure tree height. For instance, the TruPulse 200X has a manufacturer’s stated laser accuracy of +/-4.0 cm, but if the Tree-Height Routine is used the calculated height may be significantly off if the assumptions of the Tangent Method are not met. Thus, there is a distinction between the accuracy of an instrument’s sensors, e.g. a rangefinder’s laser and tilt sensor, and the measurement methods used that engage those sensors. The methods are mathematical models that work only if their assumptions fulfilled.

#### Options for Measuring Diameter

For diameters between the ground and 1.5-2.0 m, D-Tape is the main instrument. When detailed modellings are undertaken, calipers are also used. For diameters above 1.5-2.0 m, Vortex and Bushnell monoculars with range-finding reticles are ideal tools because of their superior optics, especially the Bushnell that allows seeing the reticle markings against a dark trunk. These tools are able to measure width at a distance. They have been tested repeatedly, because significant errors in diameters aloft would be translated to the average form factor.

Regarding errors in diameter measurements, D-Tape on its diameter side is calibrated to be read reliably to about 0.10 cm. Failing to keep the tape at 90° to the trunk center line and rough bark ridges can cause variations of at least 0.51 cm. Calipers can be read reliably to +/-0.13 cm. However, positioning of the instrument on the trunk leads to variations because of bark, orientation, and how tightly the instrument is pressed against the trunk. Variations of 0.51 cm are normal due to positioning and pressure. Together the error could be roughly +/-0.6 cm, slightly worse than a D-tape. However, it must be remembered that the D-tape is measuring the perimeter of a trunk and converting it to an equivalent diameter, assuming circularity.

Verifying inherent error and accuracy when measuring diameter at a distance using a laser rangefinder and a monocular with range finding reticle is more complicated. Where visibility is good, error is +/-0.64 cm. On distant targets, accuracies of about +/-1.27 cm are possible with an experienced measurer and a very accurate distance measurer like the TruPulse 200X. If visibility is poor, error is typically within +/-1.91 cm. When a steep angle to the target is involved, a more complex formula has to be used. This can introduce additional error.

The TruPoint 300 is also used for diameter at a distance and it employs a camera. Although the manufacturer reports an error of ∼3.0 cm, for distant, less distinct targets, the accuracy range is probably around +/-5.0 cm. Therefore, at the thinnest parts of a stem, and measured at a distance, this can represent a significant error.

In extensive testing the average target width error for the Bushnell over the 61 trials was 0.74 cm. Target distances were to 56.7 m, which is within the range of distances to points on an upper trunk of a tall pine. Variable definitions covering all three formulas used with a monocular are:

W = target width

M (and M_i_) = reticle reading

D = distance to center of target where target is either flat or circular

a = angle from reticle to target in vertical plane

F = manufacturer’s reticle factor

L_1_ = distance to nearer edge of target width line

L_2_ = distance to farther edge of target width line

The most commonly used formula is:

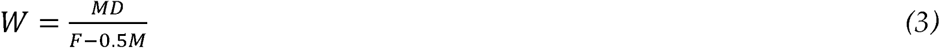

Where the target was high on a trunk the alternate formula is:

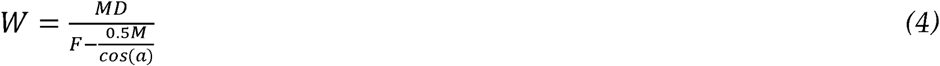

A third formula is applied to measure trunk width where there is reason to believe the trunk is not circular:

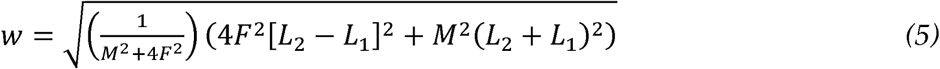

Note that the shape of the object being measured is not relevant in this third formula. It is only sensitive to the ends of the target line being measured, which can be at any orientation relative to the eye. It is the equivalent of the Missing Line routine of the TruPulse 360 and TruPoint 300.

#### Options for Measuring Trunk Volume

From the early beginnings, forest mensurationists developed statistical models for total trunk volume usually starting at 0.3 m above ground and going to total height or to a height with a 0.1 m diameter (a commercial top). Currently, the US Forest Service (USFS) uses statistically-derived trunk form or shape factors in species-specific allometric equations to compute trunk volumes. The methodology is well established, and allows volumes to be calculated simply from tree height and diameter inputs. In general, an idealized trunk’s shape changes from neiloid, to paraboloid, and finally conical, with a substantial paraboloid portion (See Figure 3).

**Figure 3.**
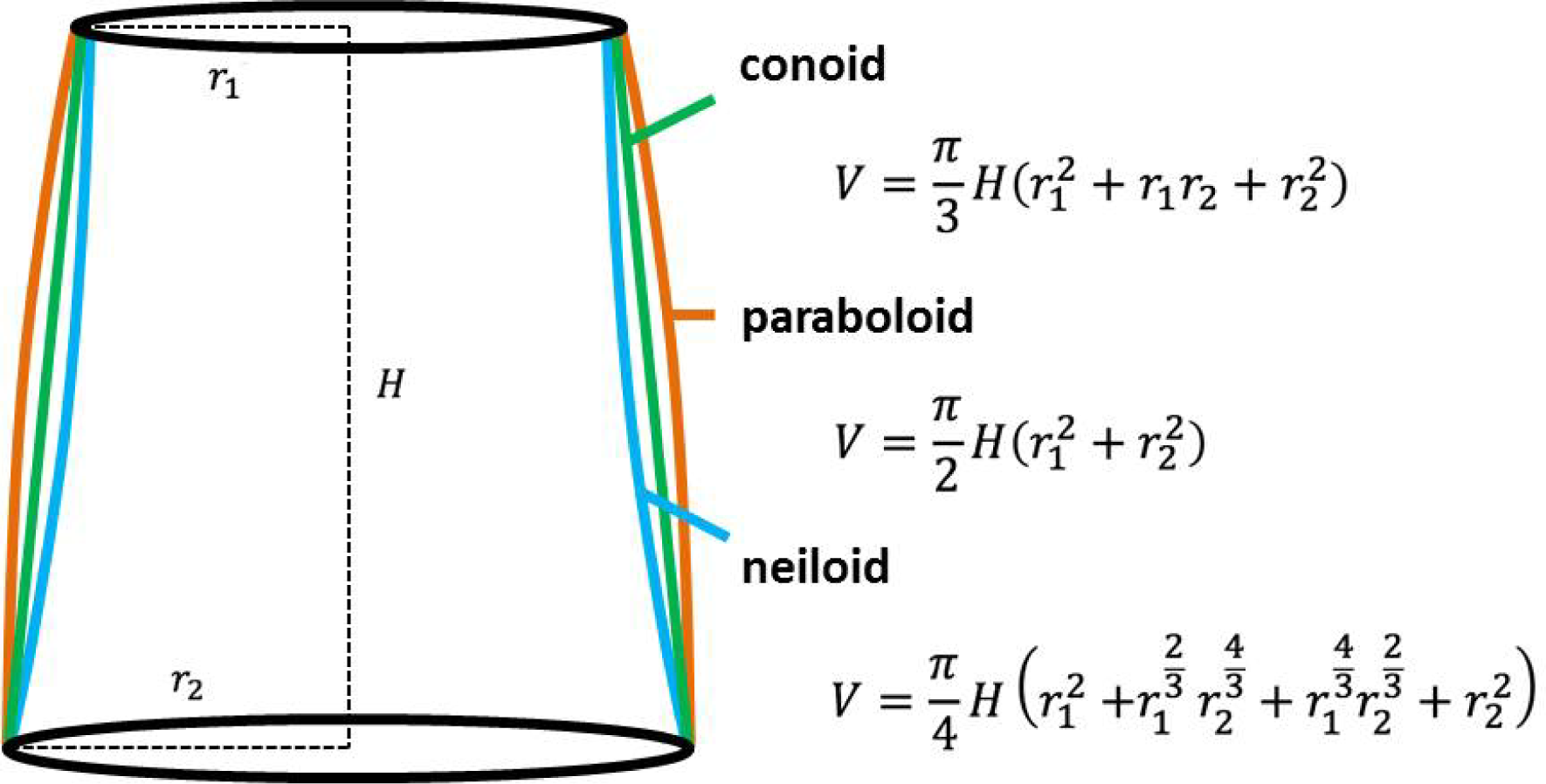
Frustums (conic or pseudo-conic forms bounded at their top by a horizontal plane parallel to the base plane) used in modeling and their defining equations. Note the convex and concave flexure of the sides of the paraboloid and neiloid, respectively.

The form factor (F in the definitions) reflects trunk shape, and can be thought of in two ways:

1. the proportion of the volume of a cylinder taken up by the trunk, the cylinder having its height equal to the height of the tree and having its base equal to the cross-sectional area of the trunk at 1.37 m above the ground. This height is adopted because it is a standard (breast height, or 4.5 feet) used in the forestry profession.
2. the average rate of trunk taper when trunk shape follows the form of a regular geometrical solid. For example, a factor of 0.333 defines a right circular cone. A form factor of 0.5 defines a quadratic paraboloid shape, and a factor of 0.25 defines a neiloid. The use of these three shapes to accurately model tree trunks is long-established [7]. Figure 3 illustrates these shapes and their corresponding equations.

#### Composite Form Factor Based on Neiloid, Cone, and Paraboloid Shapes

Geometrical solids in different combinations best model the overall shape of a trunk across the full life of a stand-grown white pine. A weighted equation for the trunk form factor (*F*) is as follows:

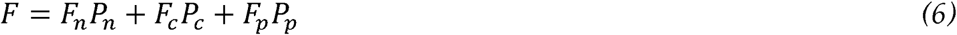

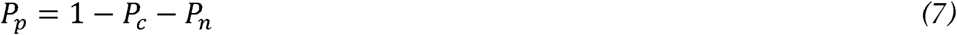

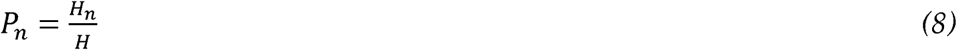

Making substitutions:

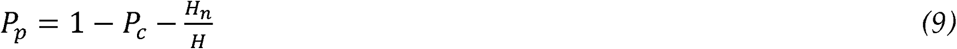

The final result is:

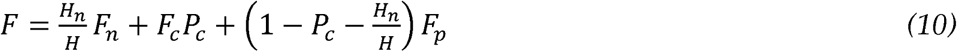

Where F_n_ =form factor for neiloid = 0.25

F_c_ = form factor for cone = 0.333

F_p_ = form factor for paraboloid = 0.5

H_n_ = length of neiloid part of trunk

H = total height of tree

P_n_ = proportion of full tree height devoted to neiloid

P_c_ = proportion of full tree height devoted to cone

P_p_ = proportion of full tree height devoted to paraboloid

#### Customized form factor based on frustums

Customized usage of multiple form factors increases the accuracy of the volume of individual trees, particularly large older trees. The customized form factor follows the contour of a series of stacked regular geometrical solids (i.e., frustums), each with one of three possible shapes (cone, neiloid, paraboloid; Fig. 3). The trunk of an individual tree is divided into geometric sections.

Where possible, diameter measurements should be made near the center of each annual internode (in the vertical direction), but higher up, foliage and limbs obscure the trunk. The form factor enters volume determinations through the following equation:

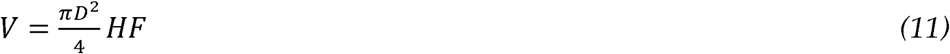

Where D = trunk diameter at breast height (1.37 m as USFS standard).

H = full tree height

F = trunk form factor

V = above ground trunk volume

Conversely to the equation above, if the volume has already been determined then the form factor is calculated as follows:

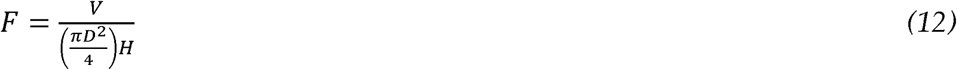

#### Limb volume

For pines grown in close proximity to one another, the common form is for a continuous trunk and lateral limbs. The trunk may be hard to discern near the top of the crown, but is easily distinguished in young trees. Since white pines put on annual branch whorls, an older tree can easily have discernible whorls with outstretched limbs for 60 or more feet.

Determining total limb volume offers challenges because of their number and visibility. In addition, white pines shed limbs from weather events over time - but that space allows existing limbs to grow larger. Detailed measures of limb structures support volumes between 16 and 21% of that of the trunk based on detailed limb measurements using a monocular with range-finding reticle, and apply to white pines grown in intact stands. The lowest value best applies to younger pines, and the range does not apply to open field-grown pines (substantially higher limb volume). Expanding further afield, hardwoods typically have a higher and species-specific limb factor. Using the form and limb factors, the trunk volume equation becomes:

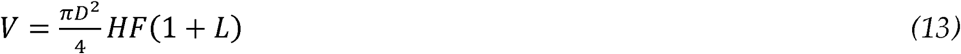

Where D = trunk diameter at breast height

H = full tree height

F = trunk form factor

L = limb factor

V = above ground trunk volume

We compare this method to a standard and commonly used USFS FIA volume-biomass model (FIAM). FIAM divides the above-ground part of a tree into stump, bole, and top (this latter includes limbs, twigs, and foliage), and here is used as implemented in the FIA worksheet BiomSlctSpp.xls [8]. FIAM uses three separate inputs: (1) stem volume is determined through a regression equation that uses diameter at breast height and height as input variables, (2) a well-known model that computes total above-ground biomass using only diameter at breast height as the input variable [9], and (3) a stump-volume model [10]. We have adjusted FIAM to express stump, bole, and top as volumes for comparability with the present method.

## Results

The customized multiple frustum method was applied to Broad Brook #2 (Figure 4), a 120-year old eastern white pine with a height typical for its age, located in a stand of other similar white pines in Massachusetts. The total height is 39.62 m, as determined by the Sine Method (*formula 2*).

**Figure 4.**
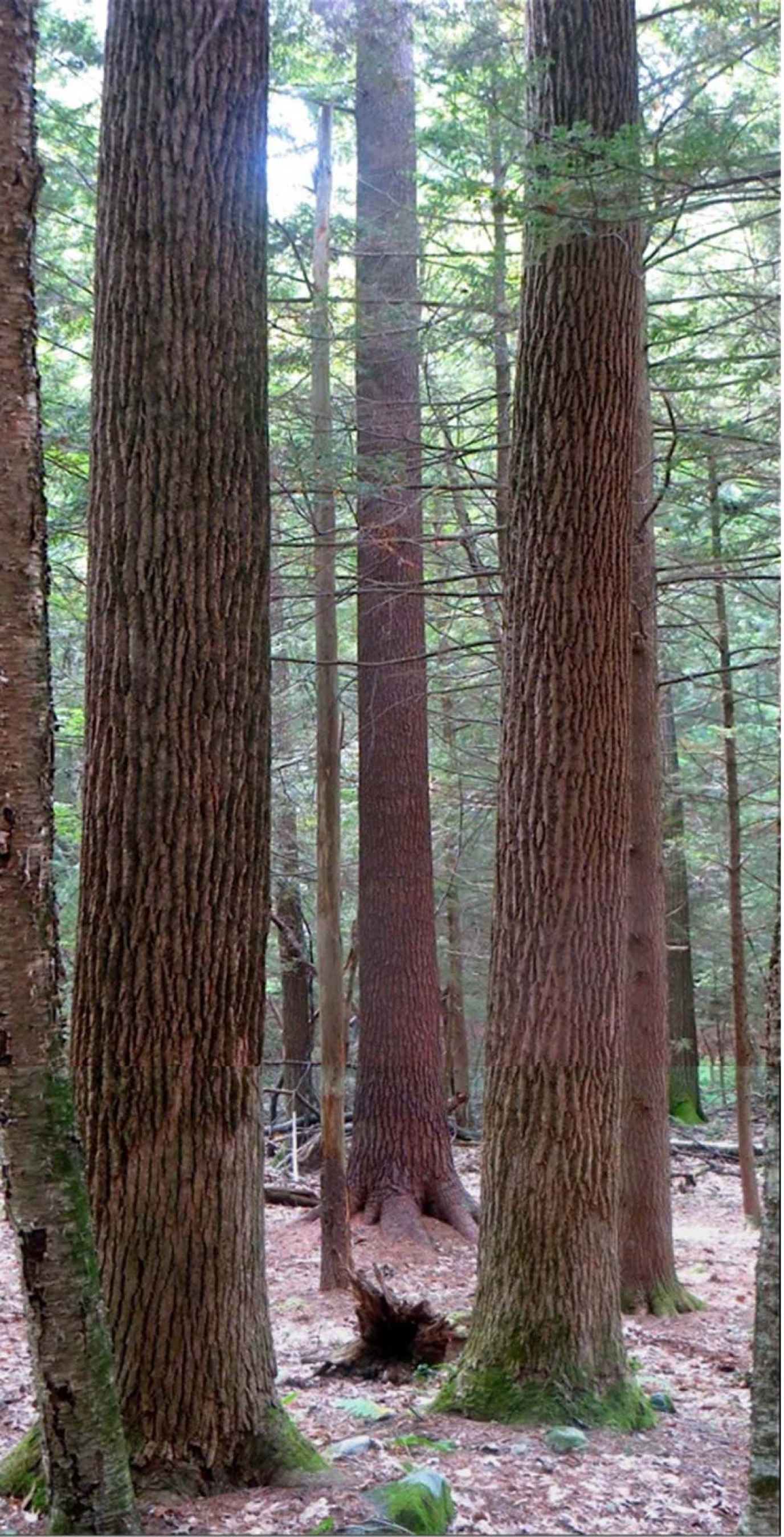
Photomontage of the lower section of a 120 y Eastern white pine in Massachusetts, catalogued as Broad Brook Pine #2 (center) seen between two foreground tuliptrees (*Liriodendron tulipifera*). The visible section of this tree is 10.7 m tall (27% of the total height), and multiple frustum modelling reveals visible above-ground volume of 3.82 m^3^ (45% of the total above-ground trunk volume). The lowest three frustums and a small part of the fourth are visible.

Individual frustums were measured (*formulas 2-4*) to generate a customized volume of the trunk of the tree; details of the frustum modeling are provided in Table 1. The trunk volume was calculated as 8.50 m^3^; back-calculating the overall form factor of the trunk (*formula 12*) gives a value of 0.405. Applying a limb factor of 15.5%, taken from a FIA-COLE model and appropriate for stand-grown white pines, leads to total above- ground volume of 9.08 m^3^ (*formula 13*) and total above-ground carbon at 1.68 tonnes (Table 1).

**Table 1:**
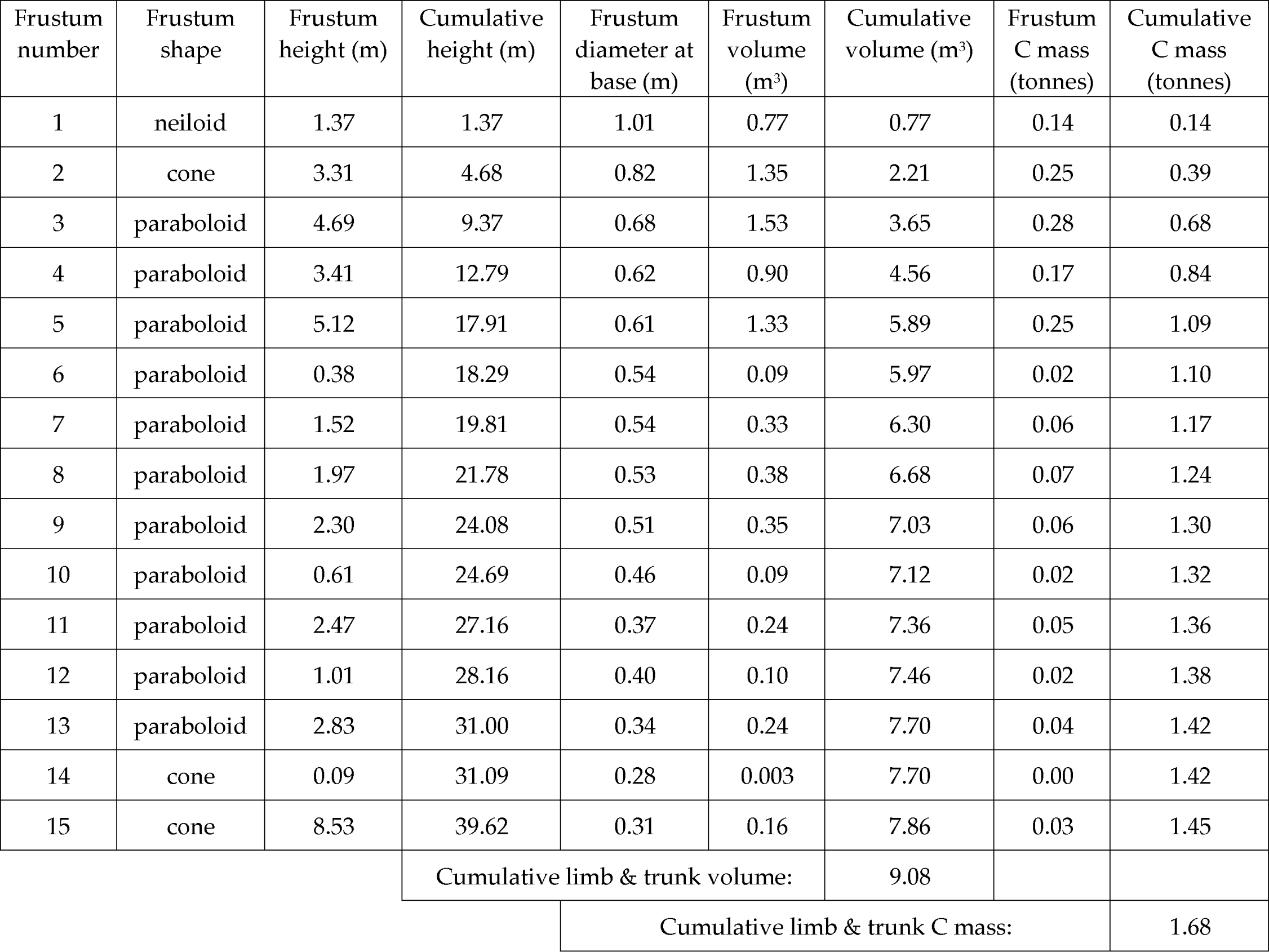
Shape modelling and carbon mass of Broad Brook Pine #2. The density of pine wood is 385.3 tonnes/m^3^ [18]. The diameter at the top of the 15^th^ frustum was considered to be 0.03 m. Conversion from volume to mass takes into account differing densities of wood and bark (bark volume is 14% of bole volume for white pine). A limb factor of 15.5% of trunk mass is applied to the carbon (C) masses. C mass is assumed to be 48% of dry wood mass. Height measurements: TruPulse 200x. Diameter measurements: Bushnell Legend Ultra HD Reticle.

This direct measurement method is compared to the FIAM statistical model for forty other white pines (Table 2) and the results are illustrated in Figure 5. Note that growth of all but one of these trees has been influenced by neighboring trees. Trunk volumes from the two methods are strongly correlated (Figure 5A) as expected with a Pearson correlation coefficient r = 0.992 (p < 0.0001). FIAM volumes are a mean 98.2% of the multiple frustum method volumes, although the two methods are not significantly different overall (Wilcoxon signed rank test Z = -1.546, p = 0.124). Differences between the two models were not related to trunk size (Figure 5B; Pearson correlation coefficient r = 0.077, p > 0.50), and FIAM is fairly accurate for most of this sample, as expected for a statistically-based model. FIAM was 15% or more off for three of these trees, however (Figure 5B). The level of accuracy of FIAM relates closely to the trunk form factor (Figure 5C; r = 0.9982):

**Table 2:**
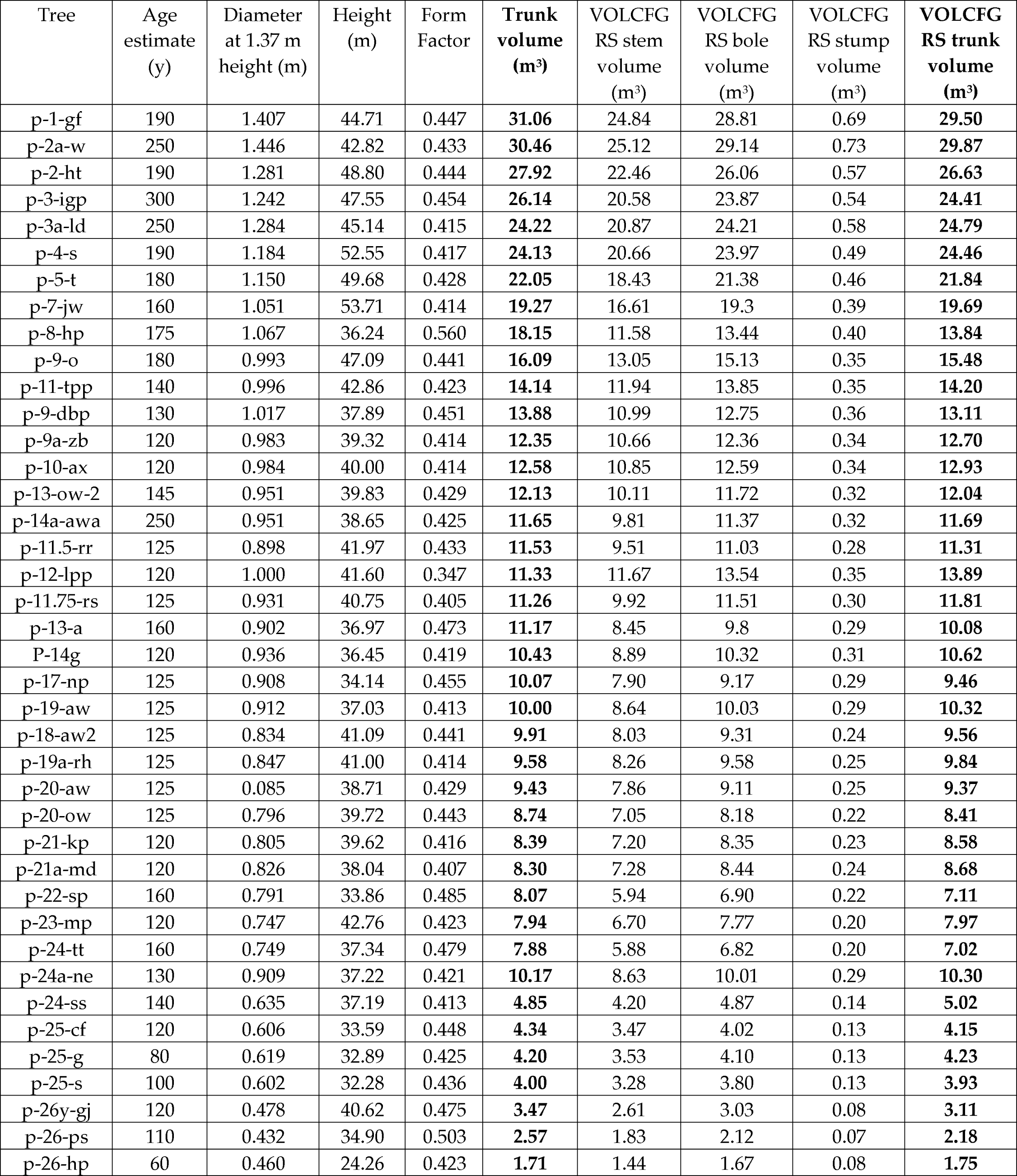
Direct measurements of other white pines and comparisons to a statistical model. Comparison of two differing tree volume measuring methods in forty white pines. The present method measures trunk volume; here, trunk is defined as bole (i.e. stem plus bark) and stump. FIA methods aim largely to measure stem volume, which represents usable lumber, but also has factors for bark and stump. The analogous factor to our “trunk” is “VOLCFGRS trunk” (volume, cubic feet gross; here converted to metric for comparability).

**Figure 5.**
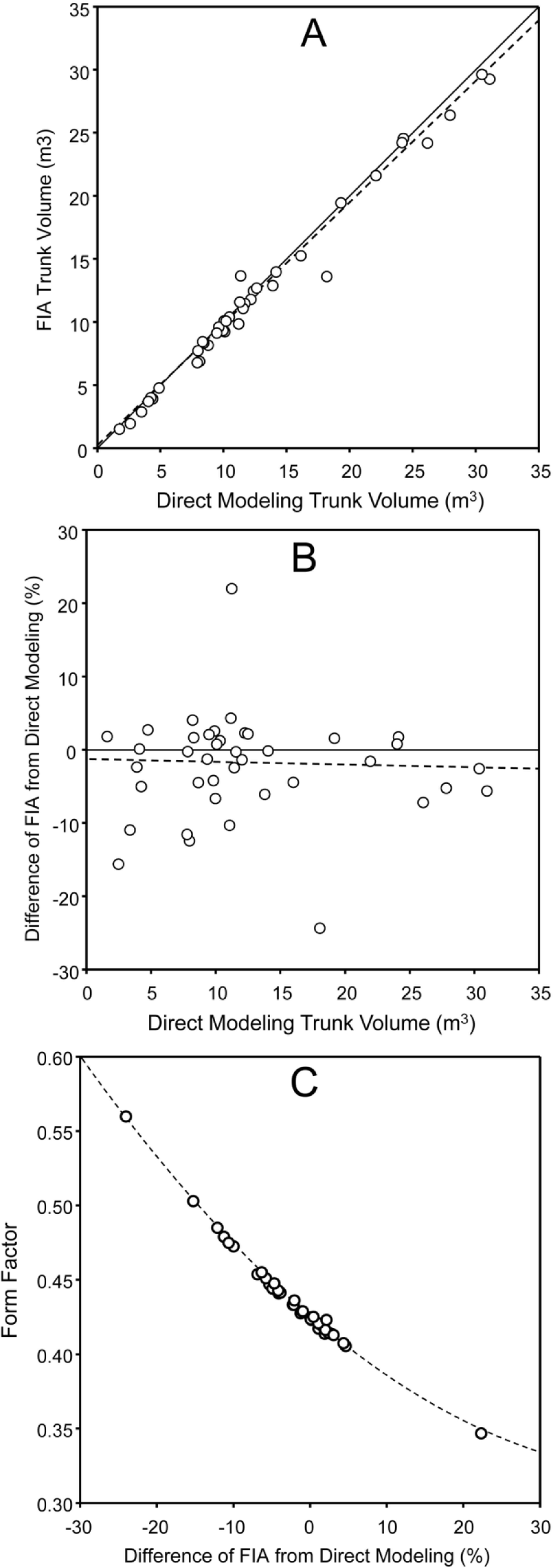
Comparison of direct modeling and FIA methods using 40 white pines. **A.** Trunk volumes from the two methods. The solid line indicates equivalence of the two models; the dashed line is the correlation of the data. **B.** Percent difference between the methods. The solid line indicates equivalence of the two models; the dashed line is the correlation of the data. **C.** Relationship of form factor to the percent difference between the methods. The dashed line is the second order regression.

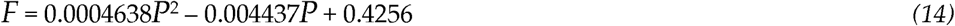

Where F = trunk form factor

P = % difference FIAM v. direct modeling

FIAM overestimates volume of the tree with the lowest form factor (P-12-LPP) by 22.6%. Conversely, FIAM underestimates volume of the trees with the two highest form factors in this sample by 23.8% (P-8-HP) and 15.0% (P-26-PS). Notably, these last two pines had previously lost their tops and had regrown bushy, wide, short tops.

## Discussion

Here we describe and apply a highly accurate direct modeling method for measuring individual trees. A white pine (120 y old) is used a detailed example, and it is applied to 40 additional trees and compared to a statistical method. The utility of the present method becomes apparent with older, larger trees that have lost apical dominance and with significant sections of parabolic trunk as well as any tree with an unusual shape.

Regarding applying this approach, it should be noted that the limb factor described applies only to stand-grown pines. The protocol is not applicable directly to species other than white pine or similar (e.g. other northeastern conifers such as red pines (*Pinus resimosa*), eastern hemlock (*Tsuga canadensis*), and red spruce (*Picea rubens*)). Furthermore, these volume determinations are limited to above-ground wood volume – they do not account for roots below ground, which add a substantial and species-specific amount to the total wood volume [11,12], nor do they account for foliage. More work needs to be done in to refine estimates for below-ground tree volume and soil carbon for specific ages and species and forest conditions.

Improvements of the present calculations could involve adaptation to multiple-trunk trees; more sophisticated treatment of non-circular trunks; and basal wedge measurement for trees on slopes. Some methods in the testing stage include frustums with elliptical cross-sections. Future advances in measuring equipment applied to the present methodology could involve unmanned aerial vehicles (e.g. [13,14], terrestrial laser scanning (e.g. [15,16]) and smartphones (apps already exist for the Tangent Method [17]).

We compared the present direct measurement method with a commonly used statistically-based forestry method for a set of forty white pines. Statistical tests indicated no overall significant difference between the two methods. At the level of individual trees, FIAM trunk volumes were quite close to those with the direct method for most of these trees, as expected for a model designed for an “average” white pine. FIAM trunk volumes were inaccurate for white pines with atypically thin or thick trunks, notably underestimating parabolic trunks. Comparisons to numerous other volume modeling methods are planned.

As forests continue to recover from historical deforestation trees will continue to get larger. Accurate individual measurements are needed to extrapolate to stands of trees or to a larger forested landscape. Because tree volume correlates directly to carbon storage, and increases over time, better quantification of the role trees and forests play in storing and sequestering carbon is urgent. This highly accurate detailed protocol can assist in targeted validation and recalibration of other standards and models that estimate volume and carbon storage over time.

## Author Contributions

Conceptualization, R.T.L; methodology, R.T.L; validation, R.T.L,, D.N.R., S.A.M.; resources, R.T.L.; data curation, D.N.R.; writing—original draft preparation, R.T.L., D.N.R., S.A.M.; writing— review and editing, R.T.L, D.N.R, S.A.M; visualization, R.T.L., D.N.R; funding acquisition, S.A.M.

## Acknowledgments

Jared D. Lockwood assisted in measuring white pines in several locations. His contribution was invaluable.

